# Cellular and molecular mechanism for reproductive capacity of male Mongolian cattle basing on single-cell sequencing for testis

**DOI:** 10.1101/2025.04.01.646534

**Authors:** Shuaipeng Gao, Shuyin Zhang, Hongrui Ren, Haoyu Yang, Li Wang, Jinxin He, Junzhen Zhang, Bayier Mengke, Lixian Zhu, Ge Qiqi, Sarengaowa Aierqing, Ruiwen Fan, Muren Herrid

## Abstract

Mongolian cattle is an excellent local breed with strong reproductive ability in Inner Mongolia of China. Spermatogenesis is a comprehensive process to decide the reproductive ability. In order to understand the cellular and molecular mechanism of spermatogenesis of Mongolian cattle, scRNA-seq was used to describe the heterogeneity of cells and gene expression profile in testis. The results showed 8 cell clusters. Among somatic cells, Sertoli cells were obviously more in young cattle than that in adult cattle, moreover, there were 4 subpopulations of Sertoli cells (SC1∼4). In the matured Sertoli cells (SC4), up-regulated genes were mainly enriched in immune system activity, enhancing immune response, and regulating cellular processes. The top 5 marker genes (*BTG2, FOSB, EGR1, JUNB, CCNL1*) were highly expressed in the testes of Mongolian cattle and water buffalo compared to Holstein cattle, which were highly expressed in the developing SC1. Gain/loss-of-function of *EGR1* in Sertoli cells isolated from Mongolian cattle testes positively regulated the expression of *FOS* and *JUN*. The results indicated that SC1 acted as the foundation and SC4 as the matured supportive cells and *EGR1* would promote the spermatogenesis by regulating AP-1 transcription factor member *FOS* and *JUN*to maintain strong reproductive capacity of Mongolian cattle.

## Introduction

Mongolian cattle is one of the excellent local breeds in north of China, which has strong working ability, endurance, and performance. Mongolian cattle are adapt to the harsh climate, and can survive and reproduce normally without the need for fine feeding and management.

Mongolian cattle typically reach sexual maturity at 12-18 months, and their semen quality is superior to many other breeds with high sperm motility and density^[1]^. Even in harsh environments such as cold and drought, Mongolian cattle maintain normal reproductive ability and exhibit excellent production performance and genetic stability^[2]^. The heterogeneity of developmental testis is helpful for understanding the cellular and molecular mechanism of reproductive performance of Mongolian cattle.

Mammalian testes is composed of seminiferous tubule and interstitial compartment with Sertoli cells and germ cells in the seminiferous tubule and Leydig cell, muscle-like cells, and macrophages in the interstitial compartment ^[3]^, which is most important male reproductive tissue for spermatogenesis and the function of both exocrine (highly specialized gametes, sperm (SP) in epididymis) and endocrine (androgen) ^[4]^. In mammals, the process of spermatogenesis requires the support of a specialized microenvironment (i.e., niche), where Sertoli and Leydig cells are the main supportive components of niche cells^[5]^. Sertoli cells are the main regulator of microenvironment^[6]^for the development of germ cells in fetal and adult testes ^[7]^. Leydig cells secret crucial hormone androgen (mostly testosterone) for maintaining male secondary sex characteristics and promoting testis development and spermatogenesis ^[8]^.

Moreover, there exists the difference of the cell composition and proportion in the testis from individuals with different ages and species^[9-10]^. The technique of single-cell RNA sequencing (scRNA-seq) has been used to construct the cellular heterogeneity and landscapes of testis in ruminant animals including sheep^[11]^, goat ^[3,12]^(Yu et al, 2021; Chen et al, 2023), cattle ^[13]^(Gao et al, 2024), and cattle-yak ^[14]^(Mipam et al, 2023). For cellular heterogeneity of testis, there are differences in different species with 3 germ cell clusters and 9 somatic cell clusters in the cattle testes^[13]^(Gao et al. 2024), and 5 germ cell clusters and 6 somatic cell clusters in dairy goat ^[12]^(Yu et al. 2021). During the entire precisely regulated process of spermatogenesis, germ cells and somatic cells undergo self-renew and differentiation, which are dynamic because of differentially expressed genes ^[15]^(Huang et al, 2022). Therefore, the physiological integrity is essentially maintained for the production of effective sperm in testes ^[16]^ (Wanjari & Gopalakrishnan, 2024). In this research, we used scRNA-seq technology to establish expression profiles of testicular cells from young and adult Mongolian cattle, analyzed cell subpopulations, and identified marker genes. Subsequently, Sertoli cells (SC) from young Mongolian cattle were extracted for analysis of the gene function in cell proliferation. In addition, we compared scRNA-seq data of testes from 3-month-old Holstein cattle and water buffalo from the GEO database deposited in NCBI and compared with that from Mongolian cattle. The results would provide valuable and theoretical foundation for improving the reproductive performance of cattle breeds living in harsh environment.

## Results

### The scRNA sequencing of testes of Mongolian cattle

To understand the developmental mechanism of the reproductive performance of male Mongolian cattle, testes tissues from healthy Mongolian cattle at 6-month-old and 16-month-old with two individuals each were collected for scRNA-seq (Fig_1_a) and showed the different cellular heterogeneity (Fig_1_b and Supplemental_Table_S1). Histological examination showed significant differences in the morphology and composition of the testes of cattle. In the testes of young cattle, the seminiferous tubules were not fully formed and obvious layer and lumen structures were lacked, while in the testes of adult cattle, the layers and lumens of the seminiferous tubules became clear and visible (Fig_1_c). To further investigate the cellular and molecular features associated with this developmental process, single-cell RNA sequencing (scRNA seq) of testes was performed using the 10x Genomics platform with 2 technical replicates for each sample, generating eight datasets (Y1_1, Y1_2, Y2_1, Y2_2, M1_1, M1_2, M2_1, M2_2). A total of 40,834 cells were obtained through standard quality control (QC) for subsequent analysis (Supplemental_Fig_S1_a). It was found that there was a significant difference in the number of cells in testes between cattle at different ages with the high similarity between technical replicates (r>0.95) (Supplemental_Fig_S1_b).

**Fig 1.**
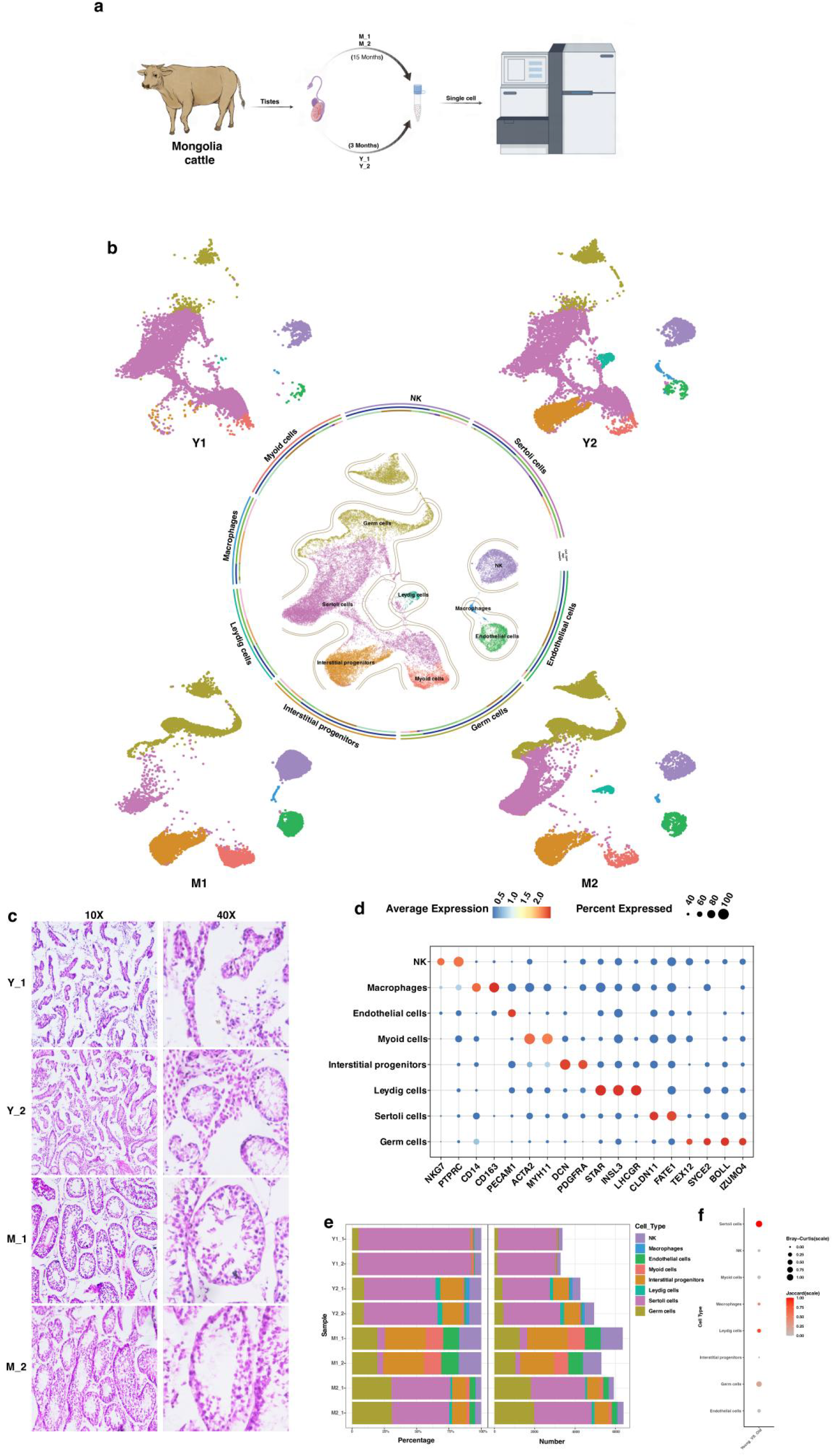
Whole information of testicular cells and marker genes expression in testis of young and adult Mongolian cattle using scRNA sequencing. a. Schematic diagram of this research. The testis tissues from healthy two 6-month-old and two 15 month old Mongolian cattle were collected for single-cell transcriptome analysis. b. UMAP maps of all testicular cells from 4 Mongolian cattle. The surroundings were UMAP maps of testicular cells from 4 individuals, different cell types shown with different colors. c. Morphological analysis of testicular tissues from individuals (paraffin slides stained by H & E, left: 10X, right: 40X). d. The marker genes used to identify cell types were displayed by dot plot. NK cells by NKG7 and PTPRC; Macrophages by CD14 and CD163; Endothelial cell by PECAM; Myoid cell by ACTA2 and MYH11; Interstitial progenitors by DCN and PDGFRA; Leydig cell by STAR, INSL3, and LHCGR; Sertoli cell by CLDN11 and FATE1; Germ cell by TEX12, SYCE2, BOLL, and IZUMO4º e. The proportion (left) and number (right) of each cell types from individuals were displayed by stacked graph. f. The difference of cells between young and adult individuals were shown by bubble chart

Using the Seurat package for Uniform Manifold Estimation and Projection (UMAP) clustering analysis, a total of 9 cell clusters were were identified and renamed according to recognized marker genes (Supplemental_Fig_S1_c). Through the analysis of the differentially expressed genes (DEGs) of each cell cluster, all somatic cell types (including NK cells, macrophages, endothelial cells, myoid cells, interstitial progenitors, leydig cells, Sertoli cells) and germ cells in the testes of Mongolian cattle were identified and labeled (Fig_1_d and Supplemental_Fig_S1_d). From the Stacked Column Diagram, the cells in testis of adult Mongolian cattle were obviously more than that in testis of young individuals. However, the proportion of Sertoli cells in testis of young Mongolian cattle was obviously higher than that in testis of adult individuals (Fig_1_e). The total of 2398 DEGs were identified (Supplemental_Fig_S1_e and Supplemental_Table_S2) and GO results were highly consistent with the biological processes of various cell types (Supplemental_Fig_S1_f). The results of Jaccard and Bray-Curtis distance showed the difference of each cell cluster in testis between young and adult Mongolian cattle, especially the obvious difference in Sertoli cells (Fig_1_f).

### The heterogeneity of Sertoli cells in testes of Mongolian cattle

To reveal the heterogeneity of Sertoli cells in testes of Mongolian cattle, all Sertoli cells from eight datasets were re-clustered and four cell subpopulation (SC_1, SC_2, SC_3, SC_4) were identified (Fig_2_a). It was showed that in testes of young individuals, the proportion of the four cell clusters was relatively uniform, however, SC4 accounted for about 70% in adult individuals (Fig_2_b). Using Monocle package, the cell trajectory found SC1 and SC4 were developing and matured Sertoli cells, respectively, while SC2 and SC3 were the transitional stages during their differentiation (Fig_2_c, Supplemental_Fig_S2_a,b). The dynamic changes in gene expression patterns at each stage were further analyzed and showed that the total number of genes, features per cell and mitochondrial genes were gradually decreased from SC1 to SC4, however, the number of mitochondrial genes were increased from SC3 to SC4 (Supplemental_Fig_S2_c). Pearson correlation analysis of different cell clusters of SC showed the highest similarity between SC3 and SC4 (R^2^ = 0.91), a significant difference between SC1 and SC3 or SC4 (R^2^ < 0.6), and SC2 in the middle (Fig_2_d). There were 501, 319, 814 and 67 DEGs in SC1∼4, respectively (Supplemental_Table_S3 and Fig_2_e). The expression of the top 5 marker genes of each cell subpopulation of SC revealed that the marker genes *BTG2, FOSB, EGR1, JUNB, and CCNL1* were top 5 in SC1, the marker genes of SC4, such as *AARD* and *FATE1*, were characteristic genes of matured Sertoli cells, and the marker genes of SC2 and SC3 were mainly related to ribosome synthesis (Fig_2_f).

**Fig 2.**
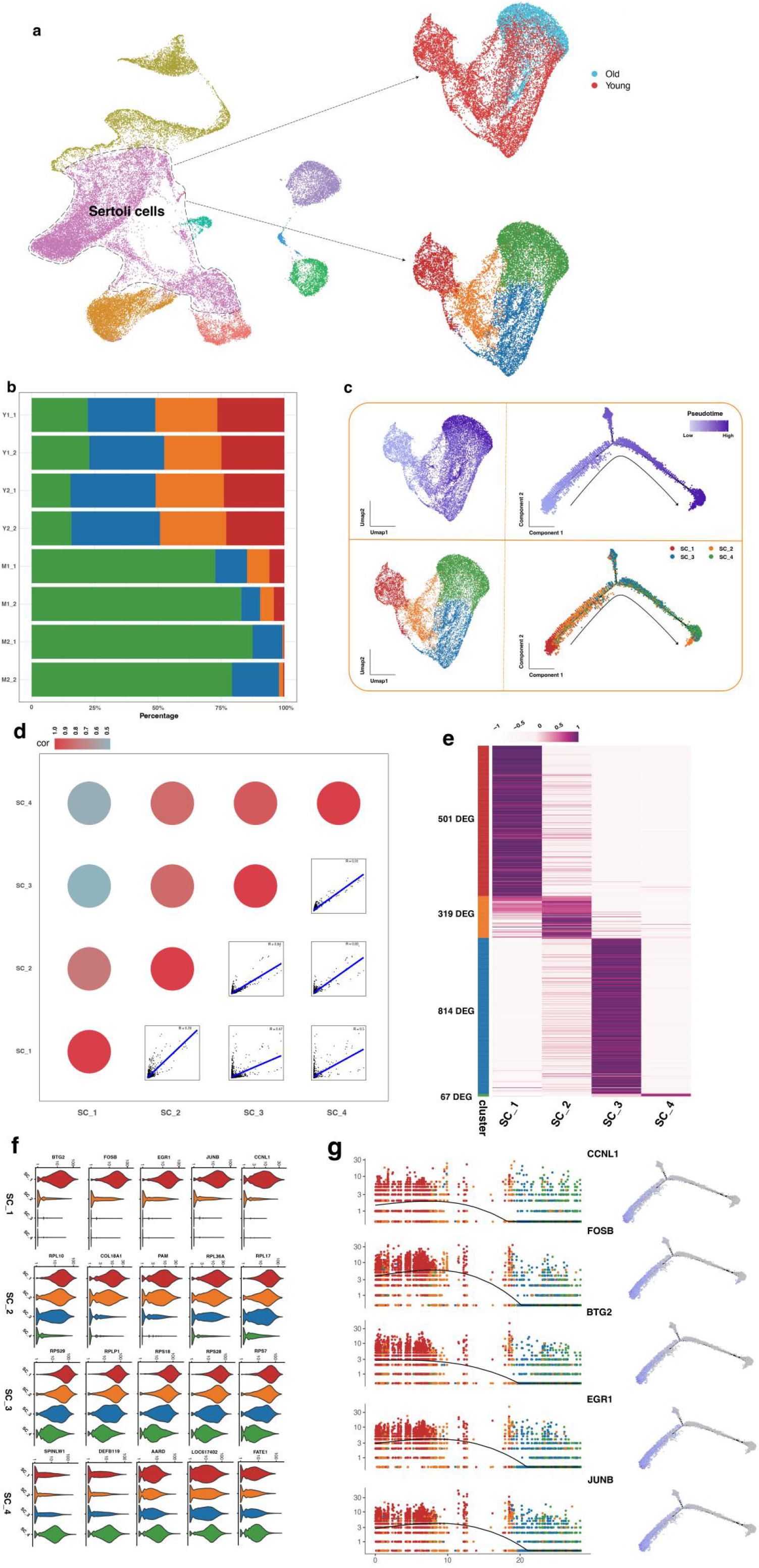
The heterogeneity of Sertoli cells in testes of Mongolian cattle. a. U-MAP for annotation of Sertoli cells re-clusters in testes from young and adult Mongolian cattle. b. The proportion of Sertoli cells subgroups for testis tissues from individuals were displayed by stacked plot with different colors (SC_1 by red; SC_2 by orange; SC_3 by blue; SC_4 by green) c. UMAP plot and pseudo time trajectory plot, where cells are stained in chronological order (top) or subpopulation (bottom). d. Pearson correlation analysis between the 4 subgroups of Sertoli cells. Bubble plot (top) with blue to red colors indicating decreasing correlation, and dot plot (bottom) with each point representing the expression of each gene in the lower and left cell subgroups. e. The heatmap of the average expression of DEGs in each subpopulation of Sertoli cells, with the count on the left side of the cell types representing the number of DEGs in each subpopulation, and the color representing the normalized average expression level of genes in cell types. f. The expression levels of the top 5 DEGs in each cell subpopulation showed by the violin plot. g. The expression patterns of the top 5 DEGs in SC-1 along the pseudo time trajectory (left) and the expression levels of these genes on the time trajectory (right).

Furthermore, the pseudo temporal analysis of the marker genes *BTG2, FOSB, EGR1, JUNB*, and *CCNL1* of SC_1 showed that the expression patterns of these 5 genes were similar. However, the expression change of *EGR1* was the most prominent with a downward trend at the earliest (Fig_2_g).

### The regulatory net in Sertoli cells of Mongolian Cattle during the development

Basing on the gene expression patterns involved in cell development trajectories, the top 200 genes with significantly dynamic expression changes were identified, including *BTG2, FOSB, EGR1*and other genes as significant initial expression genes, as well FATE1 in mature Sertoli cell as termination expression pattern genes (Fig_3_a). The correlation between these 200 genes and the trajectory Pseudotime in all Sertoli cells (Supplemental_Table_S4)and GSEA analysis for the significantly enriched functional items during the development of Mongolian Sertoli cells showed 4 pathways (Fig_3_b, Supplemental_Fig_S3_a, Supplemental_Table_S5). The MAPK signaling pathway was highly enriched in the early time series of Sertoli cells with the significant expression of *JUN, HSPA1A, HSPA8, FOS, ATF4*, and *JUN*D in SC_1 (Supplemental_Fig_S3_b). To further explore the developmental mechanism of Sertoli cells, the 455 DEGs in the SC_1 and SC_4 subpopulations with logFC greater than 2 and p < 0.05 as up-regulated genes, and 215 DEGs with logFC less than -2 and p < 0.05 as down-regulated genes were identified (Supplemental_Fig_S3_c). GO enrichment analysis showed that the up-regulated genes were mainly involved in immune system activity, enhancing immune response, regulating cellular processes, and regulating cell adhesion, while down-regulated genes were involved in the reproductive process, sperm egg binding process, and sperm production process (Fig_3_d,e). Based on the String database, PPI analysis of the highly expressed up-regulated genes in young cattle was conducted using Cytoscape, which identified the top 10 genes with the highest MCC index as hub genes, and the marker genes *EGR1, DUSP1* and *FOSB* expressed in SC1 were classified as hub genes which were connected through *FOS* and *JUN*(Fig_3_f,g).

**Fig 3.**
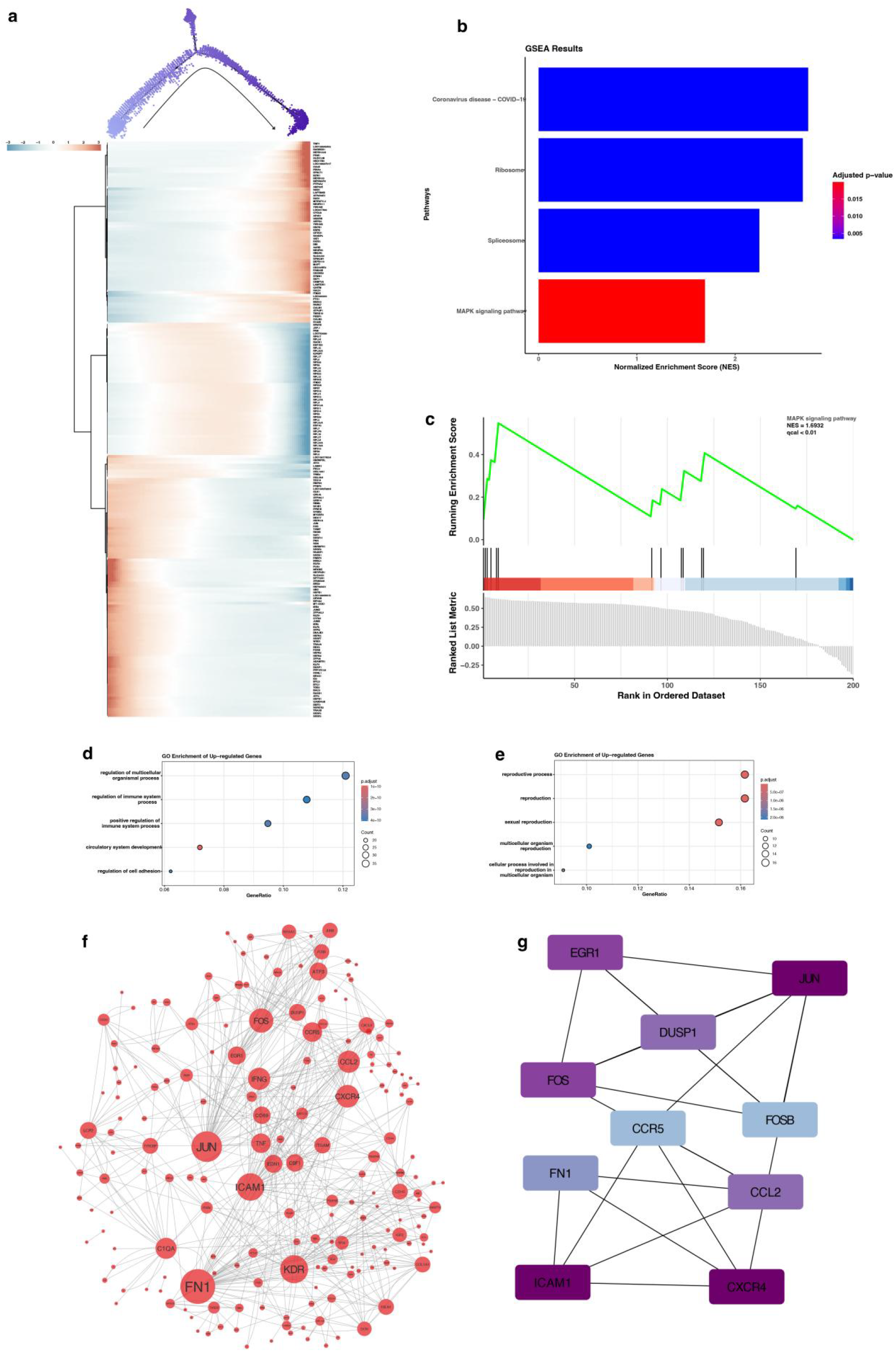
The main pathways and regulatory genes during the proliferation of Sertoli cell in testis of Mongolian cattle. a. The expression levels of the top 200 genes with significant expression differences along the pseudo time trajectory displayed by heatmap. b. The pathway and biological process of GSEA shown by a bar chart, with GSEA scores on the x-axis and color gradients changes indicating FDR values from low to high c. Schematic diagram of GSEA of MAPK pathway. d. GO enrichment of up-regulated genes shown by Bubble plot. e. GO enrichment of down-regulated genes shown by Bubble plot. f. The protein-protein interaction network of up-regulated genes with the orders of scores based on the betweenness centrality (BC). g. The top 10 genes identified by the Maximum Cluster Centrality (MCC) index using CytoHubba, with the depth of color corresponding to the weighted score.

### The comparison of Sertoli cells of Mongolian cattle with those of Holstein and Water Buffalo

In order to investigate whether SC1 acted as the foundation for Mongolian cattle to maintain strong reproductive ability in harsh environments, the scRNA-seq of testes of Mongolian cattle was compared with sequencing data of Holstein and water buffalo in public databases (water buffalo: https://www.ncbi.nlm.nih.gov/geo/query/acc.cgi?acc=GSE190477; Holstein: https://www.ncbi.nlm.nih.gov/geo/query/acc.cgi). Cell type identification was performed on these data using the Seurat package, and the expression profiles of related genes in Sertoli cells were analyzed using the marker genes (Fig_4_a and Supplemental_Fig_S4_a,b,c). The results showed the heterogeneity among samples from animals with different ages and the similarity among samples with the same age (Fig_4_c). Subsequently, Sertoli cells were re-clustered into four categories (0, 1, 2, 3) (Figure 4b). Based on the cell subclusters of SC from Mongolian cattle with different ages (Figure 4c, S. Fig. 4d), cluster 1 was determined to be the young Sertoli cell population (named as SC1). Furthermore, the expression levels of top 5 marker genes (*BTG2, FOSB, EGR1, JUNB, CCNL1*) and genes enriched in MAPK pathway showed that these gene expression patterns were highly similar with high expression in the young cell population of SC, while almost no expression in the matured SC population. At the same time, the expression of these genes was higher in the testes of Mongolian cattle and water buffalo compared to Holstein cattle (Fig_4_d).

**Fig 4.**
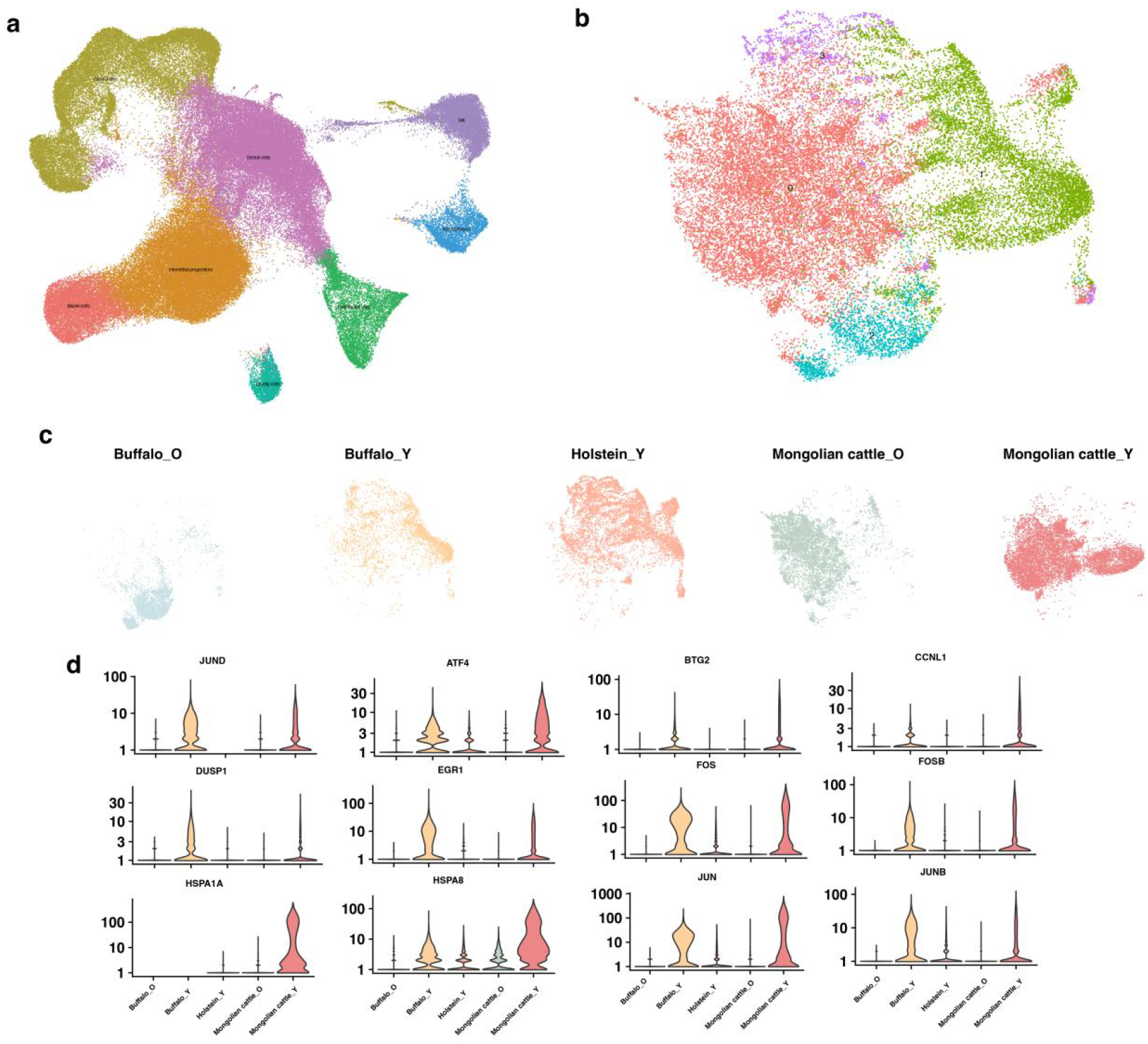
Comparison of the transcriptome analysis of single cells in testis of Mongolian Cattle,Water Buffalo, and Holstein Cattle. a. U-MAP plot for cell clusters merging the snRNA seq data of testis tissues from buffalo and Holstein cattle downloaded from NCBI. b. U-MAP plot for cell subpopulation of Sertoli cells. c. U-MAP distribution map of Sertoli cells clustered by age and species for all samples d. The expression patterns of MAPK pathway genes and key genes in all testis samples from different ages and species cattle displayed by the violin plot.

### The functional role of *EGR1*in the regulation of *JUN*and *FOS* in Sertoli cells of Mongolian cattle

To verify the expression of some DEGs, the distribution of *AMH, EGR1, FOS*and *JUN*in testes of Mongolian cattle was conducted by immunohistochemistry and the results showed that those genes were distributed in Spermatogenic epithelium of seminiferous tubules with stronger positive signals in tubules of young cattle than that of adult cattle (Fig_5_a). From the epithelium of seminiferous tubules, the Sertoli cells from Mongolian cattle testes were extracted and cultured *in vitro* and then were identified by *GATA4* and *AMH* using immunofluorescence. The co-localization of *EGR1*and *FOS*, expressed with *FOS* and *JUN*(Fig_5_b).

**Fig 5.**
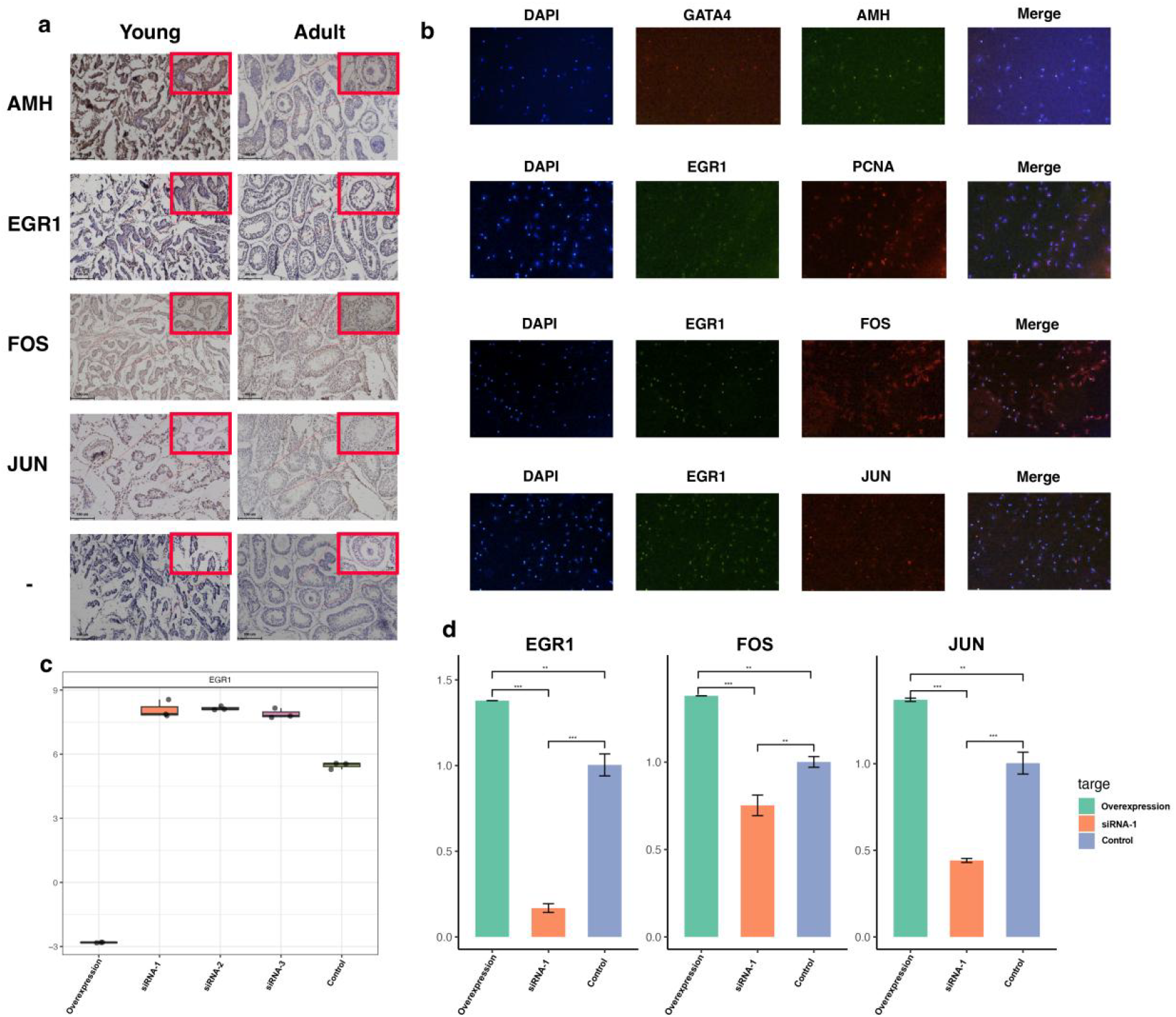
Candidate genes expression in testes tissues and SC and the regulatory role of *EGR1* in SC of Mongolian Cattle. *a*. *AMH, EGR1, FOS* and *JUN* distribution in testes of young and adult Mongolian cattle by immunohistochemistry method. b. The co-expression of *GATA* and *AMH, EGR1* and *PCNA, EGR1* and *FOS, EGR1* and *JUN* in SC of Mongolian cattle. c. the effect of 3 siRNAs and overexpression plasmid on the mRNA expression of *EGR1* d. The regulation of *EGR1* on the expression of relative genes in SC of young and adult Mongolian cattle.

Subsequently, three types of siRNA and overexpression vectors for *EGR1*were designed to transfect SC. After transfection, the total RNA of SC was extracted and performed reverse transcription to obtain cDNA, and then verified the knockout and overexpression effects of *EGR1* through qPCR. The results showed that all three siRNA could effectively downregulate the expression of *EGR1*, while overexpression vectors significantly increased the expression level of *EGR1*(Fig_5_c). Furthermore, we selected the most effective siRNA1 of *EGR1*to transfect SC, and the results showed that the mRNA expression of *JUN*and *FOS* was decreased compared to negative control and increased compared to the overexpression group with significant difference (P < 0.05) (Fig_5_d).

## Discussion

Mongolian cattle is an important local breed for milk and meat production. Though Mongolian cattle live in harsh environment, they display the better reproductive performance. To better understand structural characteristics and interaction between somatic cells and germ cells in testes of Mongolian cattle, 10 × genomics scRNA-seq was performed to investigate the cell heterogeneity and gene expression, and further comparison with Holstein cattle and water buffalo for explaining the mechanism of reproductive performance.

The lifelong fertility of male animals depends on continuous spermatogenesis, which requires the interaction between germ cells and somatic cells^[17]^ (O’donnell et al, 2022). Spermatogenesis is a comprehensive biological process with interaction of various cell types under the regulation of numerous factors, including hormones, paracrine signals, and epigenetic regulators ^[18]^(Potter & DeFalco, 2017), and is a highly complex biological process, including mitosis of spermatogonia, meiosis of spermatocytes and spermiogenesis ^[19]^(Law & Oatley, 2020). Spermatogonia includes undifferentiated type A spermatogonia (stem cell and proliferating spermatogonia) and type B differentiating spermatogonia ^[20]^(Russel et al, 1990). In pubertal testes of mammals, the differentiated spermatogonia B is developed into the primary spermatocytes by meiosis and then matured into spermatozoa ^[21]^(Bhattacharya et al, 2019). However, in testes of Mongolian cattle, 8 stages of spermagogenesis including prospermatogonia (Prospg), spermatogonial stem cells (SSCs), SPG, SPC, ES, unknown cells, RS and Sperm) were identified here. Prospg is the precursor of entire spermatogenic lineage and can be divided into three stages (mitotically active Prospg^M^, quiescent Prospg^T1^, and Prospg^T2^ after birth) ^[19]^ (Law & Oatley, 2020). Among those cells, we identified more Prospg^T2^ in testes of Mongolian cattle. Prospg^T2^ supply the resource for transiting to SSC. SSCs interact directly with Sertoli cells which are the only somatic cell type in the tubules, to control the proliferation and differentiation of cells through the secretion of specific factors ^[22]^(Chen & Liu, 2015). Therefore, Prospg^T2^ is the cellular basis for the good reproductive performance of Mongolian cattle after birth. In addition to germ cell proliferation and germ cell loss, the total output of germ cell numbers is determined by testicular size and potentially by the duration of the spermatogenic process ^[23]^(Luetjens et al, 2005). The total cells in adult Mongolian cattle testis were more than that in young cattle, which supplied the assurance for the reproductive performance.

The maturation of somatic cells is critical for the process of spermatogenesis ^[24]^(Guo et al, 2018). During early postnatal and pubertal development, Sertoli cells exhibit remarkable structural and functional change, which is necessary for the initiation and maintenance of spermatogenesis ^[25]^(Gondos & Berndston, 1994). In young Mongolian cattle, the proportion of Sertoli cells was obviously higher than that in adult cattle, which might supply the support for sprematogenesis. In pubertal cattle-yak and yak, Sertoli and Leydig cells were the main components of niche cells, while much more peritubular and perivascular cells were identified in cattle-yak^[14]^ (Mipam et al, 2023). In pubertal cattle-yak, more than 95% of the testicular cells have been identified to be niche cells, and about half of the testicular cells were niche cells in yak^[14]^ (Mipam et al, 2023). The results of yak can be attributed to the harsh living and nutritional environments of yak on Qinghai-Tibet plateau and developmental delay compared to other large domestic animals fed in lower altitude regions^[26]^. The similar proportion of Sertoli cells in testis of Mongolian cattle with yak might contribute the similar male reproductive performance under the harsh living and nutritional environment.

Moreover, 4 subpopulation of Sertoli cells were identified in Mongolian cattle, among which SC1 and SC4 were developing and matured Sertoli cells, respectively. For the comparison of gene expression between SC1 and SC4, the up-regulated genes were mainly enriched in immune system activity, enhancing immune response, regulating cellular processes and cell adhesion in SC1, while the up-regulated genes were enriched in the reproductive process, sperm binding process, and sperm production process in SC4, which supported the function of SC1 and SC4. Matured Sertoli cells not only provide nutritional and structural support but also play a major role in regulating germ cell differentiation by providing microenvironmental cues^[27]^ (Zhang et al, 2022). The expression of the marker genes of SC2 and SC3 were mainly related to ribosome synthesis, indicating that these two clusters of cells might be at the early developmental stage and had strong protein synthesis capacity, indicating that SC4 represented functionally matured Sertoli cells.

The trajectory through the marker genes expression of *BTG2, FOSB, EGR1, JUN*B, and *CCNL1*in SC1 showed that the expression trends of these 5 genes are similar with the decreased expression levels almost at the same time. However, *EGR1* showed the earliest downward trend and most prominent expression changes, indicating that *EGR1* might be a key factor regulating the expression of the associated genes. DM domain (a novel DNA-binding motif) protein (Dmrt1) is expressed in Sertoli cells and germ cells and has a direct impact on the differentiation of Sertoli cell differentiation^[28]^(Raymond et al, 2000), in which *EGR1* binds the important regulatory elements within Dmrt1 promoter to control the transcription of Dmrt1^[29]^ (Lei, 2002). DEGs were mainly enriched in the Notch, TGF-β, and Hippo signaling pathways and the signaling pathway involved in the regulation of stem cell pluripotency^[12]^ (Yu et al, 2021). From the comparison of gene expression in puberty Stertoli cells in Holstein cattle, Mongolian cattle and water buffalo, *EGR1*were highly expressed in that of Mongolian cattle and water buffalo, and low or absent in that of Holstein cattle.

The marker gene of Sertoli cells, anti-Müllerian hormone (*AMH*), acts as a direct marker of their function and indirect markers of spermatogenesis, which is a glycoprotein of the transforming growth factor-β superfamily ^[30]^(Toulis et al, 2010). The expression of *AMH*in young testes was higher than that in adult testes of Mongolian cattle, which suggested that *AMH* was produced mostly by SC in young testes and would be served as a marker for prediction of reproductive ability of young Mongolian cattle by serum *AMH*level. *GATA4*that is essential for Sertoli cell function ^[31]^(Yamamuro et al, 2021). The co-expression of *AMH* and *GATA4* in Sertoli cells suggested that the isolated cells were SC. To verify the regulatory function of *EGR1*, gain-of-function and loss-of-function of *EGR1*in Sertoli cells were conducted, which showed that *EGR1*could positively regulated *FOS* and *JUN. FOS*/*JUN*heterodimeric complex compose AP-1 transcription factor to promote the nectin-2 transcription by binding to the CRE motif ^[32]^(Lui et al, 2010). Nectin-2, a major protein component of the adherens *JUN*ction between Sertoli cells and germ cells^[32]^ (Lui et al, 2010). Also, AP-1 is involved in spermatogenesis through the regulation of androgen receptor (AR) and follicle stimulating hormone receptor (FSHR) in Sertoli cells ^[33]^(Shobana et al, 2020). Therefore, the expression of *EGR1*in Sertoli cells would promote the spermatogenesis through AP-1.

## Materials and Methods

### Animals and sample collection

The healthy Mongolian cattle (two 6-month-old and two 16 month old) were commercially obtained, which were managed on the same farm in the Alxa region of China. Cattle were cared and treated as Chi’s describe ^[34]^.

Cattle testis tissues were collected and treated as the corresponding experiments. For scRNA-seq, cells with vitality exceeding 80% were identified and captured for library constructions (10X Genomics) using Chromium Single Cell 30 Reagent Kit v3 and sequencing on the NovaSeq 6000 platform, which were as described for adipose tissue of Mongolian cattle for scRNA-seq. For immunohistochemistry and Hematoxylin & Eosin staining, the tissues were stored in 4% formaldehyde.

### The computational analysis of scRNA-seq

### The generation of raw gene extression matrix for aggregation

the version and website of the bovine reference genome and genome annotation files from NCBI GCF002263795.2-ARS-UCD1.3 (https://www.ftp.ncbi.nlm.nih.gov/genomes/all/GCF/002/263/795/GCF_002263795.2_ARS-UCD1.3/GCF_002263795.2_ARS-UCD1.3_genomic.fna.gz; https://www.ftp.ncbi.nlm.nih.gov/genomes/all/GCF/002/263/795/GCF_002263795.2_ARS-UCD1.3/GCF_002263795.2_ARS-UCD1.3_genomic.gtf.gz) were used for bioinformatics analysis. The raw scRNA-seq data was processed in Cell Ranger 6.1.2 (10x Genomics) to generate the raw gene expression matrix for aggregation using CellRanger (v5.0.1) (https://www.support.10xgenomics.com/single-cell-gene-expression/software/downloads/latest) ^[35]^. The raw sequence data reported in this paper have been deposited in the Genome Sequence Archive ^[36]^(Genomics, Proteomics & Bioinformatics 2021) in National Genomics Data Center ^[37]^(Nucleic Acids Res 2022), China National Center for Bioinformation / Beijing Institute of Genomics, Chinese Academy of Sciences (GSA: CRA024225:M1_1/M1_2/M2_1/M2_2/Y1_1/Y1_2/Y2_1/Y2_2) that are publicly accessible at https://ngdc.cncb.ac.cn/gsa.

### The quality control

The scRNA-seq of Holstein ^[38]^(GSE 244321) and Water Buffalo^[39]^ (GSE 190477) were downloaded to be analyzed together with that of Mongolian cattle generated above. Using the Read10x function to import the gene expression data into the Seurat^[40]^ (v2.3.0) R package, the cells (transcripts/cell < 200 or > 7000, > 25% mitochondrial genes) were filtered. For each sample, gene expression was expressed as a score of the gene multiplied by 10000, which was converted to natural logarithm and normalized after adding 1 to avoid taking the logarithm of 0. The top 1000 highly variable genes (HVGs) were generated from the normalized expression matrix for principal component analysis (PCA).

### Cell recognition and cluster analysis

Using Jackstrand analysis and visualization heatmaps, the significant principal components (PC) were determined especially with PC 1 to 40. Based on the local neighborhoods of all these cells, the all cells were clustered using the FindClusters (resolution = 0.4). The first 16 PCs were used to perform nonlinear dimensionality reduction through UMAP and showed as a dimensionality reduction graph. For further subclusters of Sertoli cells, FindClusters (resolution = 0.1) was performed and the first 8 PC were used for nonlinear dimensionality reduction through UMAP.

### Identification of the differentially expressed genes (DEGs)

The specific marker genes of the unique cluster were identified using the Seurat FindAllMarkers function (test. Use = wilcox) based on standardized Unique Molecular Identifier (UMI) counts. Only genes expressed in at least 25 % of cells were tested with the average log 2 (fold change) threshold set as 2 in the analysis of cell types. The DEGs in each sub-cluster were calculated based on comparisons between one of sub-clusters and the others. GO analysis was performed using WebGestalt 2019^[41]^.

### Analysis of differences between cell clusters in testes of young and adult Mongolian cattle

Based on cell clusters and sample types, these cells were grouped into sub-clusters using Seurat. Then, the average expression of the top 1000 DEGs in each sub-cluster was calculated and scaled to balance the differences in overall expression levels between sub-clusters. Afterwards, the Jaccard and Bray Curtis distances between every two sub-clusters were calculated using the Vegan ^[42]^2.5.6 R package based on the scaling values, which were displayed for the same cell types between young and adult testis as a bubble plot.

### Cell trajectory analysis

The single-cell pseudo time trajectory was constructed using Monocle 2 software package (version 2.8.0) (http://cole-trapnell-lab.github.io/monocle-release/docs_mobile/). Briefly, the UMI count matrix of Sertoli cells as expr_matrix and meta.data were input and then 1725 DEGs between clusters were used to define the progression of the cell, among which the genes expressed in less than 10 cells with P-value greater than 0.001 were excluded (eliminating batch effects by setting residualmodelFormulaStr parameter). Cells were ordered using the orderCells after reduction of the space to two dimensions using DDRTree.

### Gene Set Enrichment Analysis (GSEA) analysis

Based on the identification of genes significantly correlated with pseudo-time based on pseudo-time trajectories, the top 200 genes with the highest correlation were used to calculate the Spearman correlation coefficients between the gene expression levels and pseudo-time and order the genes according to their correlation. The Gene Set Enrichment Analysis (GSEA) method was used to perform functional enrichment analysis to reveal biological pathways and molecular functions associated with pseudo-time.

### Construction of PPI Network and Identification of Hub Genes

The specific genes of subclusters of Sertoli cells were identified using FindAllMarkers analysis in Seurat and limma packages to screen DEGs with logFC > 2 and P value < 0.05. The interactions between target DEGs were predicted by construction of protein-protein interaction (PPI) network using the STRING database (version 11.5, https://string-db.org/)with an interaction score 0.4 ^[43]^. Then, the hub genes were screened and network visualization was performed using Cytoscape software (version 3.10.0) ^[44]^. The cluster function modules in PPI networks were constructed using the Molecular Complex Detection (MCODE) plugin, and then, the nodes of target network were ordered using the Maximum Cluster Centrality (MCC) method in CytoHubba plugin basing on the values degree, betweenness centrality, and centroid of each gene within the network calculated by CentiScaPe plugin.

### Hematoxylin & Eosin (H & E) staining of testis

The fresh testis tissues of the cattle were fixed in 4% paraformaldehyde at room temperature for 18 hours, embedded in paraffin, and sliced it into 6 um slides. Before staining, the tissue slices were dewaxed in xylene, rehydrated using gradient concentrations of ethanol, and washed in distilled water. Then slides were stained with hematoxylin and eosin, which were sealed by neutral resin after dehydration with increasing concentrations of ethanol and xylene and then take stained images using microscope (Nikon).

### Immunohistochemistry

The paraffin sections of testes tissues from young and adult Mongolian cattle were dewaxed and rehydrated. The primary antibody *AMH, EGR1, FOS*and *JUN* were incubated at various concentrations, respectively, IgG as the negative control. Then the tissues were incubated with HRP-labeled anti-rabbit/mouse IgG antibody (1:50, Sigma) for 30 min, and then analyzed under microscope for *AMH, EGR1, FOS*and *JUN* distribution and quantitative expression.

### Isolation, Culture, and Identification of Primary Sertoli Cell

After washing fresh testes tissue with cold PBS containing penicillin and streptomycin, 3cm^3^ of cortex was taken and cut into small pieces. Cells were isolated from the seminiferous tubules according to the following method: the testicular tissue was digested with a mixed solution containing 2.5mg/ml trypsin EDTA and 10ug/ml DNase I, and incubated by shaking at 100 rmp at 37 °C for 20 minutes, and then trypsin activity was neutralized in a basic culture medium containing 10% FBS. Following the release of interstitial cells in PBS at room temperature for 10 minutes, the supernatant was discarded and the tubules were incubated with PBS containing 1mg/ml collagenase, 1mg/ml hyaluronidase, and 10ug/ml DNase I by shaking at 100 rpm at 37 °C for 15 minutes. After wash with PBS, the tubules was filtered through a 70um cell filter to remove cell aggregates, centrifuged at 200g at 4 °C to collect the pellets of Sertoli cells and germ cells. For getting pure Sertoli cells, the cells were grown in DMEM/F12 medium containing 10% FBS, penicillin, and streptomycin, supplemented with 1X ITS and 5-20ng/ml EGF in a 5% CO_2_ incubator at 37 °C and passaged every 3 days using trypsin/EDTA. After two days, cells were treated with 20mM Tris solution (pH 7.5) for 120 s to induce hypotonic shock of germ cells to get pure Sertoli cells for transfection.

### Immunofluorescence

Sertoli cells were grown on slides and identified by marker genes using the method of Immunofluorescence. Briefly, the slides were fixed by 4 % paraformaldehyde at room temperature for 20 minutes, which were incubated at room temperature with 1% Triton X-100 in PBS for 20 minutes. After block at room temperature with 3% bovine serum albumin (BSA) in PBS for 1 hour, the primary antibodies against *EGR1*(dilution 1:100, rabbit resource, Bioss), *FOS* (dilution 1:100, mouse resource, Bioss), *JUN*(dilution 1:100, mouse resource, Bioss), *GATA4* (dilution 1:100, mouse resource, Boster), *AMH* (dilution 1:100, rabbit resource, Bioss) were added to incubate overnight at 4 °C, respectively. Then, the secondary antibody (anti-rabbit FITC from goat, Bioss; anti-mouse 594 from goat, Proteintech) was added to incubate at room temperature in the dark for 30 minutes for being sealed by Anti fluorescence attenuation sealing agent with DAPI (Solarbio, Beijing, China). The fluorescence signals were captured under microscope (Nikon)

### Cell transfection and Quantitative real-time PCR (qPCR)

Three siRNAs for *EGR1* (Supplemental_Table_S6) were designed and synthesized (Sangon Biotech, Shanghai, China) and the overexpression plasmid with CDS (NM_001045875.1) region of *EGR1*(pcDNA 3.1-*EGR1*) were constructed. Sertoli cells were incubated in a 6-well plate to grow at a confluence of 50-70%, which were used to transfect with *EGR1*-siRNAs and overexprssion plasmid using Lipofectamine 2000 (Invitrogen) in DMEM/F12 medium with antibiotic free according to the manufacture’s instructions. The experiment was divided into *EGR1*-siRNA1 group, *EGR1*-siRNA2 group, *EGR1*-siRNA3 group, *EGR1*overexpression group, and control group (pcDNA3.1). After the cell transfection and incubation in medium at 37 °C for 6-8 hours, cells were cultured in the fresh complete medium for 24 hours. Then, cells were collected and total RNAs were extracted using Trizol reagent for qPCR. qPCR (Supplemental_Table_S7) and the 2 ^-△△ ct^ method were used to detect the expression of *EGR1, FOS* and *JUN*.

### Statistical analysis

All data from qPCR were expressed as mean with standard error of the mean (Mean ± SEM). The differences among groups were analyzed by ANOVA and Tukey HSD in rstatix^[45]^ package, and \**p*<0.05, \*\**p*<0.01, \*\*\**p*<0.001 were considered statistically significant.

## Competing Interest Statement

The authors declare that there was no competing Interests in this work.

## Acknowledgment

We thank the fund support by the projected “Revelation and Leadership” project of the Department of Science and Technology of Inner Mongolian Autonomous Region (2022JBGS0023).

## Author Contributions

S.G., R.F., M.H., designed and wrote the main manuscript. S.G., S.Z., H.R, H.Y., performed experiments. S.G, L.W., R.F., processed and analyzed the data. M.B., Z.L., Q.G., A.S., fed cattle, collected samples and supplied the formation of cattle. H.J., Z.J., reviewed and edited the manuscript.

## References

[1] Ll Xin, HAN Bi-Ying, YANG Ming, ZHANG xue-Li, HAl Chao, Ll Guang-Peng, ZHAO Yue-Fang. Correlation Analysis Between Amino Acid Content in Bovine(Bos taurusSeminal Plasma and Sperm Motility

[2] CHEN Jia-lei, XlA Xiao-ting, XlAO Zheng-zhong, CHEN Ning-bo. LEl Chu-zhac. The Genetic Diversity of Mitochondrial DNA Genome of Mongolian Cattle

[3] Chen M, Wang N, Yang H, Liu D, Gao Y, Duo L, Cui X, Hao F, Ye J, Gao F, Tu Q, Gui Y. Single-cell transcriptome analysis of the germ cells and somatic cells during mitotic quiescence stage in goats. FASEB J. 2023 Nov;37(11):e23244. doi: 10.1096/fj.202301278

[4] Törzsök P, Steiner C, Pallauf M, Abenhardt M, Milinovic L, Plank B, Rückl A, Sieberer M, Lusuardi L, Deininger S. Long-Term Follow-Up after Testicular Torsion: Prospective Evaluation of Endocrine and Exocrine Testicular Function, Fertility, Oxidative Stress and Erectile Function. J Clin Med. 2022 Nov 2;11(21):6507. doi: 10.3390/jcm11216507.

[5] Morrison SJ, Spradling AC. Stem cells and niches: mechanisms that promote stem cell maintenance throughout life. Cell. 2008 Feb 22;132(4):598–611. doi: 10.1016/j.cell.2008.01.038.

[6] Hai Y, Hou J, Liu Y, Liu Y, Yang H, Li Z, He Z. The roles and regulation of Sertoli cells in fate determinations of spermatogonial stem cells and spermatogenesis. Semin Cell Dev Biol. 2014 May;29:66–75. doi: 10.1016/j.semcdb.2014.04.007.

[7] Rebourcet D, Darbey A, Monteiro A, Soffientini U, Tsai YT, Handel I, Pitetti JL, Nef S, Smith LB, O’Shaughnessy PJ. Sertoli Cell Number Defines and Predicts Germ and Leydig Cell Population Sizes in the Adult Mouse Testis. Endocrinology. 2017 Sep 1;158(9):2955–2969. doi: 10.1210/en.2017-00196.

[8] Ramm SA, Schärer L, Ehmcke J, Wistuba J. Sperm competition and the evolution of spermatogenesis. Mol Hum Reprod. 2014 Dec;20(12):1169–79. doi: 10.1093/molehr/gau070.

[9] Hermann BP, Cheng K, Singh A, Roa-De La Cruz L, Mutoji KN, Chen IC, Gildersleeve H, Lehle JD, Mayo M, Westernströer B, Law NC, Oatley MJ, Velte EK, Niedenberger BA, Fritze D, Silber S, Geyer CB, Oatley JM, McCarrey JR. The Mammalian Spermatogenesis Single-Cell Transcriptome, from Spermatogonial Stem Cells to Spermatids. Cell Rep. 2018 Nov 6;25(6):1650-1667.e8. doi: 10.1016/j.celrep.2018.10.026.

[10] Shami AN, Zheng X, Munyoki SK, Ma Q, Manske GL, Green CD, Sukhwani M, Orwig KE, Li JZ, Hammoud SS. Single-Cell RNA Sequencing of Human, Macaque, and Mouse Testes Uncovers Conserved and Divergent Features of Mammalian Spermatogenesis. Dev Cell. 2020 Aug 24;54(4):529-547.e12. doi: 10.1016/j.devcel.2020.05.010. Epub 2020 Jun 5. PMID: 32504559; PMCID: PMC7879256.

[11] Yang H, Ma J, Wan Z, Wang Q, Wang Z, Zhao J, Wang F, Zhang Y. Characterization of sheep spermatogenesis through single-cell RNA sequencing. FASEB J. 2021 Feb;35(2):e21187. doi: 10.1096/fj.202001035RRR.

[12] Yu XW, Li TT, D. XM, Shen QY, Zhang MF, Wei YD, Yang DH, Xu WJ, Chen WB, Bai CL, Li XL, Li GP, Li N, Peng S, Liao MZ, Hua JL. Single-cell RNA sequencing reveals atlas of dairy goat testis cells. Zool Res. 2021 Jul 18;42(4):401–405. doi: 10.24272/j.issn.2095-8137.2020.373.

[13] Gao Y, Bai F, Zhang Q, et al. Dynamic transcriptome profiles and novel markers in bovine spermatogenesis revealed by single-cell sequencing[J]. Journal of Integrative Agriculture, 2024, 23(7):2362–23

[14] Mipam T, Chen X, Zhao W, Zhang P, Chai Z, Yue B, Luo H, Wang J, Wang H, Wu Z, Wang J, Wang M, Wang H, Zhang M, Wang H, Jing K, Zhong J, Cai X. Single-cell transcriptome analysis and in vitro differentiation of testicular cells reveal novel insights into male sterility of the interspecific hybrid cattle-yak. BMC Genomics. 2023 Mar 27;24(1):149. doi: 10.1186/s12864-023-09251-2.

[15] Huang L, Zhang J, Zhang P, Huang X, Yang W, Liu R, Sun Q, Lu Y, Zhang M, Fu Q. Single-cell RNA sequencing uncovers dynamic roadmap and cell-cell communication during buffalo spermatogenesis. iScience. 2022 Dec 6;26(1):105733. doi: 10.1016/j.isci.2022.105733.

[16] Wanjari UR, Gopalakrishnan AV. Blood-testis barrier: a review on regulators in maintaining cell junction integrity between Sertoli cells. Cell Tissue Res. 2024 May;396(2):157–175. doi: 10.1007/s00441-024-03894-7.

[17] O’Donnell L, Smith LB, Rebourcet D. Sertoli cells as key drivers of testis function. Semin Cell Dev Biol. 2022 Jan;121:2–9. doi: 10.1016/j.semcdb.2021.06.016.

[18] Potter SJ, DeFalco T. Role of the testis interstitial compartment in spermatogonial stem cell function. Reproduction. 2017 Apr;153(4):R151–R162. doi: 10.1530/REP-16-0588.

[19] Law NC, Oatley JM. Developmental underpinnings of spermatogonial stem cell establishment. Andrology. 2020 Jul;8(4):852–861. doi: 10.1111/andr.12810.

[20] Russel, L. D., Ettlin, R. A., Sinha Hikim, A.P., Clegg, E. D. (1990). Histological and histopathological evaluation of the testis. Cache River Press, Clearwater. Florida.

[21] Bhattacharya I, Sen Sharma S, Majumdar SS. Pubertal orchestration of hormones and testis in primates. Mol Reprod Dev. 2019 Nov;86(11):1505–1530. doi: 10.1002/mrd.23246.

[22] Chen SR, Liu YX. Regulation of spermatogonial stem cell self-renewal and spermatocyte meiosis by Sertoli cell signaling. Reproduction. 2015 Apr;149(4):R159–67. doi: 10.1530/REP-14-0481.

[23] Luetjens CM, Weinbauer GF, Wistuba J. Primate spermatogenesis: new insights into comparative testicular organisation, spermatogenic efficiency and endocrine control. Biol Rev Camb Philos Soc. 2005 Aug;80(3):475–88. doi: 10.1017/s1464793105006755.

[24] Guo J, Grow EJ, Mlcochova H, Maher GJ, Lindskog C, Nie X, Guo Y, Takei Y, Yun J, Cai L, Kim R, Carrell DT, Goriely A, Hotaling JM, Cairns BR. The adult human testis transcriptional cell atlas. Cell Res. 2018 Dec;28(12):1141–1157. doi: 10.1038/s41422-018-0099-2.

[25] Gondos B, Berndston WE. Postnatal and pubertal development. In: Knobil E, Neill JD (eds.), The Physiology of Reproduction, 2nd ed. New York: Raven Press; 1994: 116–154.

[26] Wang G, Li Y, Yang Q, Xu S, Ma S, Yan R, Zhang R, Jia G, Ai D, Yang Q. Gene expression dynamics during the gonocyte to spermatogonia transition and spermatogenesis in the domestic yak. J Anim Sci Biotechnol. 2019 Jul 12;10:64. doi: 10.1186/s40104-019-0360-7.

[27] Zhang L, Guo M, Liu Z, Liu R, Zheng Y, Yu T, Lv Y, Lu H, Zeng W, Zhang T, Pan C. Single-cell RNA-seq analysis of testicular somatic cell development in pigs. J Genet Genomics. 2022 Nov;49(11):1016–1028. doi: 10.1016/j.jgg.2022.03.014.

[28] Raymond CS, Murphy MW, O’Sullivan MG, Bardwell VJ, Zarkower D. Dmrt1, a gene related to worm and fly sexual regulators, is required for mammalian testis differentiation. Genes Dev. 2000 Oct 15;14(20):2587–95. doi: 10.1101/gad.834100.

[29] Lei N, Heckert LL. Sp1 and Egr1 regulate transcription of the Dmrt1 gene in Sertoli cells. Biol Reprod. 2002 Mar;66(3):675–84. doi: 10.1095/biolreprod66.3.675.

[30] Toulis KA, Iliadou PK, Venetis CA, Tsametis C, Tarlatzis BC, Papadimas I, Goulis DG. Inhibin B and anti-Mullerian hormone as markers of persistent spermatogenesis in men with non-obstructive azoospermia: a meta-analysis of diagnostic accuracy studies. Hum Reprod Update. 2010 Nov-Dec;16(6):713–24. doi: 10.1093/humupd/dmq024.

[31] Yamamuro T, Nakamura S, Yamano Y, Endo T, Yanagawa K, Tokumura A, Matsumura T, Kobayashi K, Mori H, Enokidani Y, Yoshida G, Imoto H, Kawabata T, Hamasaki M, Kuma A, Kuribayashi S, Takezawa K, Okada Y, Ozawa M, Fukuhara S, Shinohara T, Ikawa M, Yoshimori T. Rubicon prevents autophagic degradation of GATA4 to promote Sertoli cell function. PLoS Genet. 2021 Aug 5;17(8):e1009688. doi: 10.1371/journal.pgen.1009688.

[32] Lui WY, Sze KL, Lee WM. Nectin-2 expression in testicular cells is controlled via the functional cooperation between transcription factors of the Sp1, CREB, and AP-1 families. J Cell Physiol. 2006 Apr;207(1):144–57. doi: 10.1002/jcp.20545.

[33] Shobana N, Kumar MK, Navin AK, Akbarsha MA, Aruldhas MM. Prenatal exposure to excess chromium attenuates transcription factors regulating expression of androgen and follicle stimulating hormone receptors in Sertoli cells of prepuberal rats. Chem Biol Interact. 2020 Sep 1;328:109188. doi: 10.1016/j.cbi.2020.109188.

[34] Chi Z, Jia Q, Yang H, Ren H, Jin C, He J, Wuri N, Sui Z, Zhang J, Mengke B, Zhu L, Qiqi G, Aierqing S, Wuli J, Ai D, Fan R, Herrid M. snRNA-seq of adipose tissues reveals the potential cellular and molecular mechanisms of cold and disease resistance in Mongolian cattle. BMC Genomics. 2024 Oct 25;25(1):999. doi: 10.1186/s12864-024-10913-y. PMID: 39448899; PMCID: PMC11520132.

[35] Butler A, Hoffman P, Smibert P, Papalexi E, Satija R. Integrating single-cell transcriptomic data across different conditions, technologies, and species. Nat Biotechnol. 2018 Jun;36(5):411–420. doi: 10.1038/nbt.4096.

[36] Chen T, Chen X, Zhang S, Zhu J, Tang B, Wang A, Dong L, Zhang Z, Yu C, Sun Y, Chi L, Chen H, Zhai S, Sun Y, Lan L, Zhang X, Xiao J, Bao Y, Wang Y, Zhang Z, Zhao W. The Genome Sequence Archive Family: Toward Explosive Data Growth and Diverse Data Types. Genomics Proteomics Bioinformatics. 2021 Aug;19(4):578–583. doi: 10.1016/j.gpb.2021.08.001.

[37] CNCB-NGDC Members and Partners. Database Resources of the National Genomics Data Center, China National Center for Bioinformation in 2022. Nucleic Acids Res. 2022 Jan 7;50(D1):D27-D38. doi: 10.1093/nar/gkab951.

[38] Mueller ML, McNabb BR, Owen JR, Hennig SL, Ledesma AV, Angove ML, Conley AJ, Ross PJ, Van Eenennaam AL. Germline ablation achieved via CRISPR/Cas9 targeting of NANOS3 in bovine zygotes. Front Genome Ed. 2023 Nov 27;5:1321243. doi: 10.3389/fgeed.2023.1321243.

[39] Huang L, Zhang J, Zhang P, Huang X, Yang W, Liu R, Sun Q, Lu Y, Zhang M, Fu Q. Single-cell RNA sequencing uncovers dynamic roadmap and cell-cell communication during buffalo spermatogenesis. iScience. 2022 Dec 6;26(1):105733. doi: 10.1016/j.isci.2022.105733.

[40] Satija R, Farrell JA, Gennert D, Schier AF, Regev A. Spatial reconstruction of single-cell gene expression data. Nat Biotechnol. 2015 May;33(5):495–502. doi: 10.1038/nbt.3192.

[41] Xu S, Hu E, Cai Y, Xie Z, Luo X, Zhan L, Tang W, Wang Q, Liu B, Wang R, Xie W, Wu T, Xie L, Yu G. Using clusterProfiler to characterize multiomics data. Nat Protoc. 2024 Nov;19(11):3292–3320. doi: 10.1038/s41596-024-01020-z.

[42] Oksanen, Jari, et al. Vegan: Community Ecology Package. Version 2.6-8, 2024, R package.

[43] Szklarczyk D, Morris JH, Cook H, Kuhn M, Wyder S, Simonovic M, Santos A, Doncheva NT, Roth A, Bork P, Jensen LJ, von Mering C. The STRING database in 2017: quality-controlled protein-protein association networks, made broadly accessible. Nucleic Acids Res. 2017 Jan 4;45(D1):D362–D368. doi: 10.1093/nar/gkw937.

[44] Shannon P, Markiel A, Ozier O, Baliga NS, Wang JT, Ramage D, Amin N, Schwikowski B, Ideker T. Cytoscape: a software environment for integrated models of biomolecular interaction networks. Genome Res. 2003 Nov;13(11):2498–504. doi: 10.1101/gr.1239303.

[45] Kassambara, Alboukadel. Rstatix: Pipe-Friendly Framework for Basic Statistical Tests. Version 0.7.2, 2023, R package

